# Dihydromyricetin alleviated the damage of hypoxia-induced mouse neurons by reducing ROS levels and inhibiting the expression of PAR and γH2AX

**DOI:** 10.1101/2024.07.07.602423

**Authors:** Xueping Du, Yanjun Guo, Junzheng Yang

**Author notes:** Corresponding author: Junzheng Yang, Guangdong Nephrotic Drug Engineering Technology Research Center, The R&D Center of Drug for Renal Diseases, Consun Pharmaceutical Group, Guangzhou 510530, China.

## Abstract

**Objective:** To investigate the effect of dihydromyricetin on hypoxia-induced neurons, to understand the effect of dihydromyricetin on hypoxic-ischemic encephalopathy (HIE).

**Methods:** Cortical neurons were isolated from C57BL/6j mice (24 hour-year old), cultured, and subjected to 4h hypoxia and 20h reoxygenation to mimic the neonatal hypoxic-ischemic encephalopathy. After dihydromyricetin (20μmol/L) treatment of hypoxia-induced neurons for 2h, CCK-8 assay was used to analyze the neuronal viability, Hoechst33342/PI double staining assay was used to analyze the neuronal death, Western blotting was used to analyze the expression of Poly ADP-ribose (PAR) polymer protein and _γ_H2AX, comet assay was used to detect DNA damage, immunofluorescence staining was used to observe the nuclear translocation of apoptosis inducing factor, and 2’,7’-dichlorodihydrofluorescein diacetate was used to detect the expression of reactive oxygen species (ROS).

**Results:** Compared with the control groups, hypoxia-treated neurons exhibited significantly lower activity, higher neuronal death rate and the high expressions of PAR and _γ_H2AX, hypoxia could also induce AIF nuclear translocation, increase tail DNA content and tail length, increase the expression of ROS in neurons; after dihydromyricetin treatment, neuronal activity were significantly increased, neuronal death rate, ROS levels, and the expressions of PAR and _γ_H2AX were also decreased, AIF nuclear translocation was inhibited, the tail DNA content and tail length were also decreased.

**Conclusion:** Dihydromyricetin could alleviate the damage of hypoxia-induced neurons through decreasing the levels of ROS and inhibiting the expressions of PAR and _γ_H2AX, suggesting that dihydromyricetin may have the protective effect on HIE.

## 1 Introduction

Neonatal hypoxic-ischemic encephalopathy (HIE) is a common cause of neonatal death and long-term neurological dysfunction, it could result from reduced cerebral blood flow and hypoxic-ischemic brain injury due to various causes during the perinatal period [1, 2]. It is one of the most common causes of postnatal disability in newborns [3]. It is reported that perinatal asphyxia is the main cause of HIE, any risk factors which could cause disturbances in blood circulation and gas exchange between the mother and fetus, will result in a decrease in blood oxygen concentration, can cause asphyxia, and further lead to HIE [4, 5]. The statistical results demonstrated that the incidence rate of HIE in United Kingdom in 2021 was 2.96‰ [6], the incidence rate in developing countries reached up to 26‰ [7], especially, the risk of adverse outcomes for surviving newborns can reach up to 50% [8]; almost of 50% HIE patients were caused by intrauterine asphyxia, almost 40% of HIE patients were caused by asphyxia during delivery, and 10% of HIE patients were caused by congenital diseases [9, 10].

There is a lack of the ideal treatment for HIE, the current treatment methods include correction of hypoxaemia and hypercapnia [11], correction of hypotension [12], control of convulsions by sodium phenobarbital [13], control of increased intracranial pressure by dexamethasone [14], central nervous system stimulants [15], but there were many application limitations of their treatment methods. Dihydromyricetin is widely present in plants of the grape family and genus Ophiopogon, it has various pharmacological activities including anti-tumor effect, antioxidant effect, anti-apoptotic effect, and anti-inflammatory effect [16, 17, 18]. However, whether dihydromyricetin has the protective effect on HIE was not clear. In this article, we treated the hypoxia-induced mouse neurons with dihydromyricetin, and discussed the underlying mechanisms, hoping to provide some useful clues for application of dihydromyricetin in HIE.

## 2 Materials and Methods

### 2.1 Main reagents

Dihydromyricetin and dimethyl sulfoxide (DMSO) were purchased from Shanghai Yuanye Biotechnology Co., Ltd; CCK-8 kit, Hoechst33342/PI kit, and ROS kit were purchased from Shanghai Biyuntian Biotechnology Co., Ltd; comet electrophoresis kit was purchased from Shanghai Kaiji Biotechnology Co., Ltd; AIF antibody, PAR antibody, _γ_H2AX antibody and actin antibody were purchased from CST company, Alexa Fluor 594 secondary antibody, goat anti-rabbit HRP IgG, and anti-fluorescence quencher (including DAPI) were purchased from Beijing Zhongshan Jinqiao Biotechnology Co., Ltd. Neurobasal culture medium, L-glutamine, and B-27 were purchased from Invitrogen company.

### 2.2 Isolation and culture of cortical neurons, and the establishment of the hypoxia-induced neuron model

Mice were purchased from Huaxing Experimental Animal Farm (license number was SYXK (Yu) 2020-0004). Cortical neurons were isolated from C57BL/6 24-hour-old mice and cultured in Neurobasal culture medium at 37°C in a 5% CO_2_ environment. After 7 days of culture, the normal culture medium was replaced with sugar-free culture medium, the cortical neurons were kept in a three-gas incubator (37°C, 5% CO_2_, 1% O_2_, and 94% N_2_) for 4h; and the sugar-free culture medium was replaced with normal culture medium, and the cells were further cultured in the 37°C, 5% CO_2_ incubator for another 20h; finally, 20_μ_mol/L dihydromyricetin was used to treat the hypoxia-induced neuron for 2h, and the cortical neurons were assessed by multiple detection methods.

### 2.3 Hoechst 33342/PI experiment

The treated cortical neurons were collected and resuspended in 1mL culture medium, 10uL of Hoechst staining solution was added and incubated in the dark room (37°C, 10 min), centrifuged at 800g for 5min, and the supernatant was aspirated, 1mL working solution and 5uL PI staining solution were added to resuspend the neurons in the dark room at the room temperature for 20min; finally, the neurons were observed and photographed using a fluorescence microscope.

### 2.4 Reactive oxygen species detection assay

The treated neurons were washed with PBS and incubated with 10μmol/L DCFH-DA for 20min in the dark; after washing with PBS, they were observed and photographed under a fluorescence microscope.

### 2.5 CCK-8 proliferation assay

The cortical neuron was seeded in the 96 well plate. After processing according to the experimental requirements, 10uL CCK-8 working solution was added into each well and incubated at 37°C for 1h; the absorbance values (450nm) of each well were measured using an enzyme linked immunosorbent assay (ELISA) reader.

### 2.6 Immunofluorescence staining

The treated neurons were fixed with 4% paraformaldehyde for 15 min at room temperature, permeabilized with TritonX-100 for 20 min, blocked with 3% bovine serum albumin (BSA) for 1h, incubated with rabbit anti-AIF primary antibody (1:150) overnight at 4°C, after washing the neurons with PBS, incubated with goat anti-rabbit secondary antibody conjugated with AlexaFluor 594 for 1h at room temperature; after washing by PBS for three times, the nuclei were re-stained with DAPI; and observed and photographed under a confocal fluorescence microscope.

### 2.7 Western blotting

Neurons were harvested before and after treatment, and protein was extracted using RIPA buffer. Protein concentration was measured by using BCA method. The denatured protein was separated by using SDS-PAGE and then transferred to a PVDF membrane. The membrane was blocked with 5% skim milk at room temperature for 1h, and the primary antibody was incubated overnight at 4°C. after washing with TBST, the corresponding secondary antibody was incubated for another 1h at 37°C. After incubation with ECL-luminescent solution, the PVDF membrane was developed and t photographed using a multifunctional imaging device.

### 2.8 Comet assay

The treated neurons were washed and resuspended with the pre-cooled PBS at the concentration of 1×10^6^/mL; the slides were taken out and pre-coated with 1% agarose (normal dissolution point), and then the second layer of cell agarose solution (including resuspended cells and 0.7% agarose (the ratio was 1:7.5 (v/v) with low melting point) was added, the slides were kept in the dark room at 4°C for 30min to increase adhesion; the slides covered with 0.7% agarose with low melting point and stored at 4°C until solidified; then the cells on the slides were lysed in the pre-cooled lysis buffer at 4°C for 1h, and then the slides were immersed in alkaline dissociation solution at room temperature for 1h; the electrophoresis was performed in the horizontal electrophoresis device for 30min at the voltage of 21V; after rinsing with Tris-HCl (0.4mmol/L, pH 7.5) for three times, the glass slides were stained with PI at 4°C in dark rooms for 10min, observed and photographed under the fluorescence microscope, and finally the number of tail-positive cells, the DNA content (%) in the tail, and the length of the tail (from the edge of the nucleus to the end of the comet tail) were counted manually.

## 3 Statistical methods

Data conforming to the normal distribution were represented as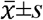 and were analyzed by using SPSS 23.0 software; t-test was used to compare the differences between two groups, one-way ANOVA was used to compare the differences among multiple groups; LSD test was used for pairwise comparison of different groups. P<0.05 was considered the significant statistical difference.

## 4 Results

### 4.1 Dihydromyricetin could increase the cell viability and decrease the apoptosis rate and ROS in the cortical neurons

Compared with the normal neurons, the viability of cortical neurons cultured under the hypoxic environment was significantly reduced, and the number of PI-positive cortical neurons (apoptotic cells) was increased (P<0.01); after dihydromyricetin treatment, the neuronal activity was increased and the PI positive cortical neurons were decreased significantly (P<0.01), and dihydromyricetin only had no effect on the cell viability and the apoptosis rate in the normal cortical neurons (Figure 1 and Table 1).

**Figure 1.**
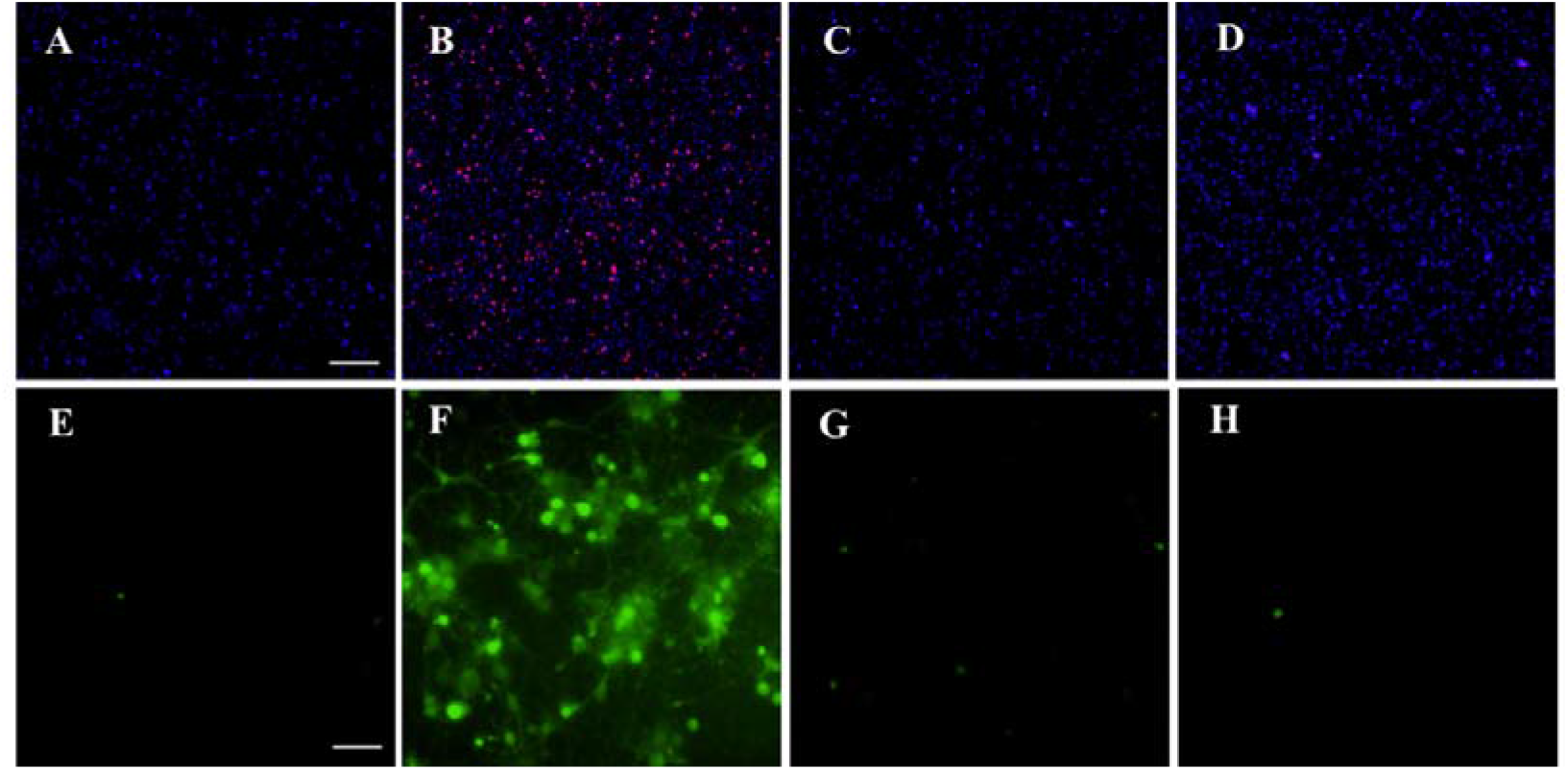
Hoechst33342/PI staining observed and the ROS experiments detected the effects of dihydromyricetin on hypoxia-induced cortical neurons. A-D: Hoechst33342/PI staining observed the effects of dihydromyricetin on hypoxia-induced cortical neurons in control group (A), OGD/R group (B), DI+ OGD/R group (C) and DI group (D), respectively; E-H: the ROS experiments detected the effects of dihydromyricetin on hypoxia-induced cortical neurons in control group (E), OGD/R group (F), DI+ OGD/R group (G) and DI group (H), respectively. Scale bar=20μm or scale bar=40μm.

**Table 1.** The changes of cell vitality, PI positive rate, and ROS of cortical neurons in different group 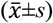.

| Group | Sample size | Vitality | PI-positive rate (%) | ROS |
| --- | --- | --- | --- | --- |
| Control group | 3 | 1.03±0.04 | 5.98±1.83 | 4.33±1.53 |
| OGD/R group | 3 | 0.55±0.06<sup>##</sup> | 43.64±8.40<sup>##</sup> | 91.67±27.54<sup>##</sup> |
| DI+ OGD/R group | 3 | 0.84±0.03<sup>**</sup> | 10.75±2.24<sup>**</sup> | 21.33±8.50<sup>**</sup> |
| DI group | 3 | 0.90±0.06 | 5.92±1.48 | 5.67±2.52 |
| *F* |  | 48.44 | 17.08 | 24.43 |
| *P* |  | <0.001 | <0.01 | <0.001 |

Abbreviations: OGD/R, hypoxia/reoxygenation; DI, dihydromyricetin; Compared with the control group, <sup>#</sup>P<0.05, <sup>##</sup>P<0.01, <sup>###</sup>P<0.001; compared with the DI+OGD/R group, <sup>\*</sup>P<0.05, <sup>\*\*</sup>P<0.01, <sup>\*\*\*</sup>P<0.001.

### 4.2 Dihydromyricetin could regulate tail length and DNA content in the tail

Compared with the normal cortical neurons, increased DNA, and length in the tail of cortical neurons were found (P<0.01); and after dihydromyricetin treatment, increased DNA, and length in the tail of cortical neurons were improved (P<0.01), and dihydromyricetin only had no effect on tail length and DNA content in the tail in the normal cortical neurons (Figure 2 and Table 2).

**Figure 2.**
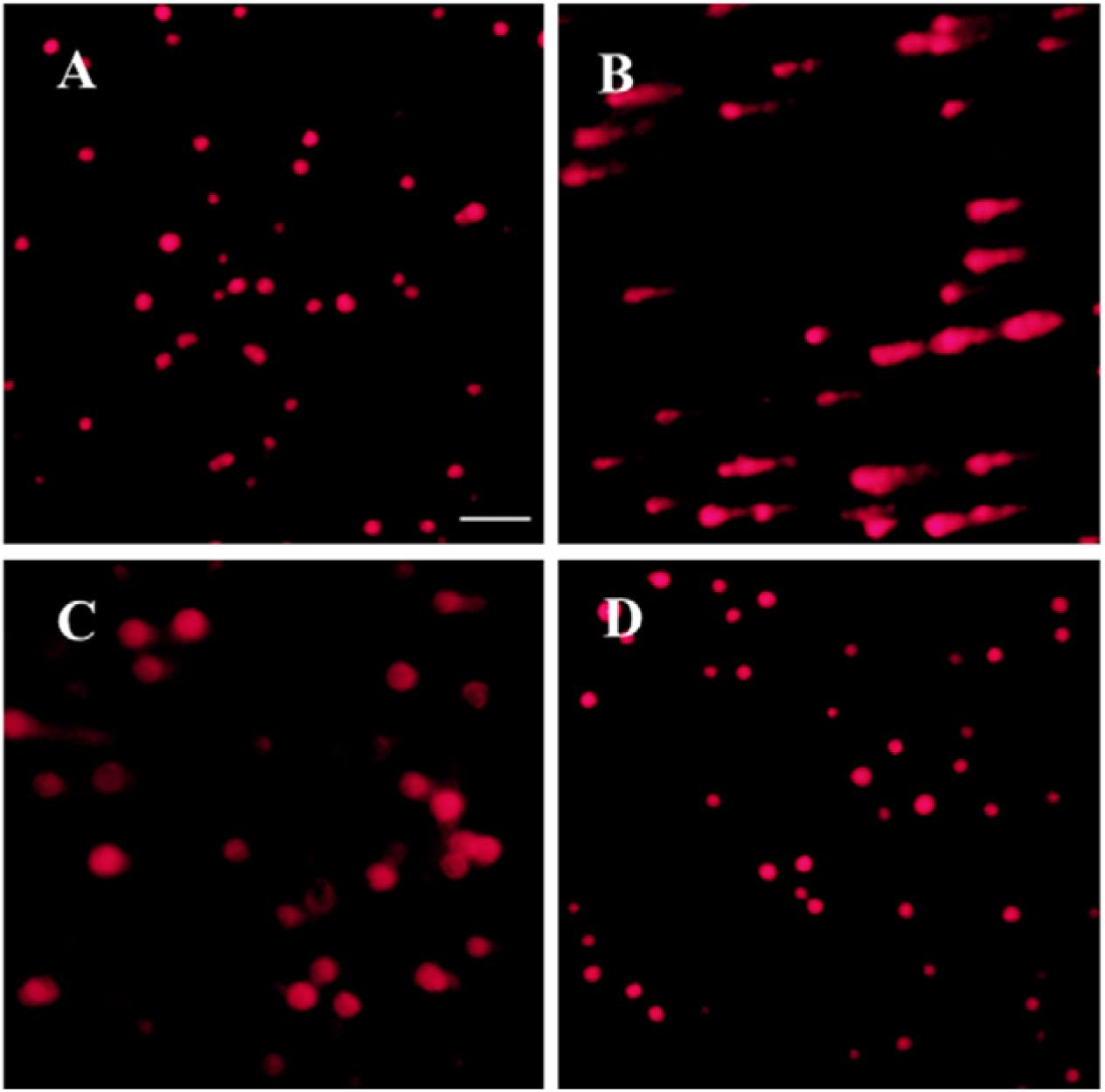
Effects of dihydromyricetin on the tail length and tail DNA content in cortical neurons. A: control group; B: OGD/R group; C: DI+ OGD/R group; D: DI group.

**Table 2.** Comparison of tail length and DNA content in the tail of cortical neurons in different groups 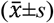.

| Group | Sample size | Tail length ($\mu\text{m}$) | DNA content in tail (%) |
| --- | --- | --- | --- |
| Control group | 3 | $6.37 \pm 1.25$ | $1.80 \pm 0.71$ |
| OGD/R group | 3 | $41.60 \pm 2.50^{##}$ | $27.39 \pm 3.71^{##}$ |
| DI+ OGD/R group | 3 | $8.71 \pm 0.26^{**}$ | $4.50 \pm 0.45^{**}$ |
| DI group | 3 | $6.57 \pm 0.80$ | $1.83 \pm 1.01$ |
| *F* |  | 418.29 | 119.23 |
| *P* |  | <0.001 | <0.001 |

Abbreviations: OGD/R, hypoxia/reoxygenation; DI, Dihydromyricetin; Compared with the control group, $^{\#}P < 0.05$ , $^{##}P < 0.001$ , $^{###}P < 0.001$ ; Compared with the DI+OGD/R group, $^{*}P < 0.05$ , $^{**}P < 0.01$ , $^{***}P < 0.001$ .

### 4.3 Dihydromyricetin could regulate the AIF nuclear translocation in cortical neurons

The immunofluorescence staining results demonstrated that the AIF nuclear translocation was found compared with the normal cortical neurons (P<0.01); and after dihydromyricetin treatment, AIF nuclear translocation of neurons was improved (P<0.01), and dihydromyricetin had no effect on AIF nuclear translocation in the normal cortical neurons (Figure 3).

**Figure 3.**
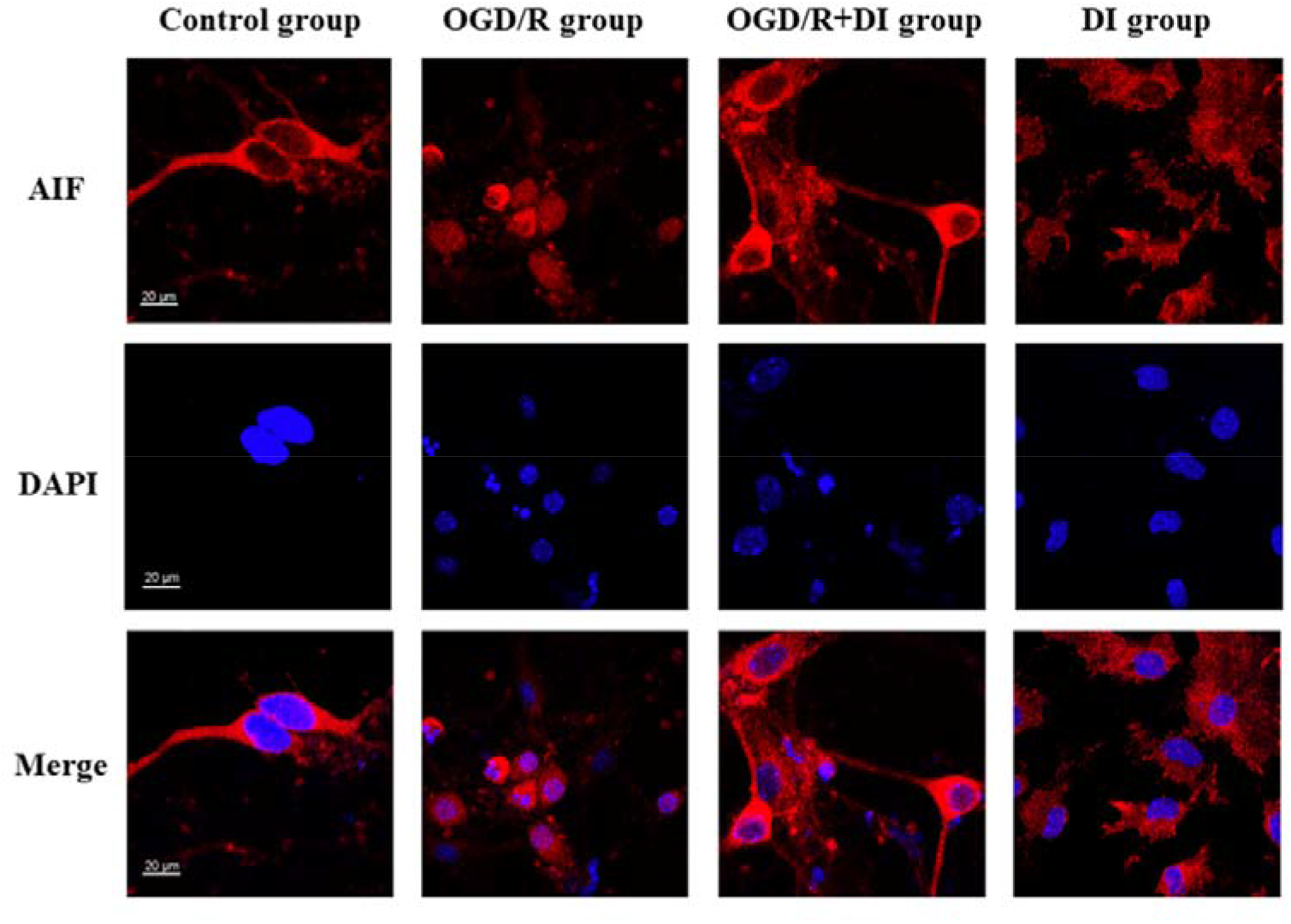
Immunofluorescence staining observed the nuclear translocation of AIF in cortical neurons in different groups. Abbreviations: AIF, apoptosis inducing factor; DAPI, 4’,6-diamidino-2-phenylindole, scale bar=20μm.

### 4.4 Dihydromyricetin could regulate the expression of PAR and _γ_H2AX in cortical neurons

Compared with the normal cortical neurons, the expressions of PAR and _γ_H2AX in the hypoxia group were increased (P<0.01); and after dihydromyricetin treatment, the expressions of PAR and _γ_H2AX were decreased (P<0.01), and dihydromyricetin only had no effect on AIF nuclear translocation in the normal cortical neurons (Figure 4 and Table 4).

**Figure 4.**
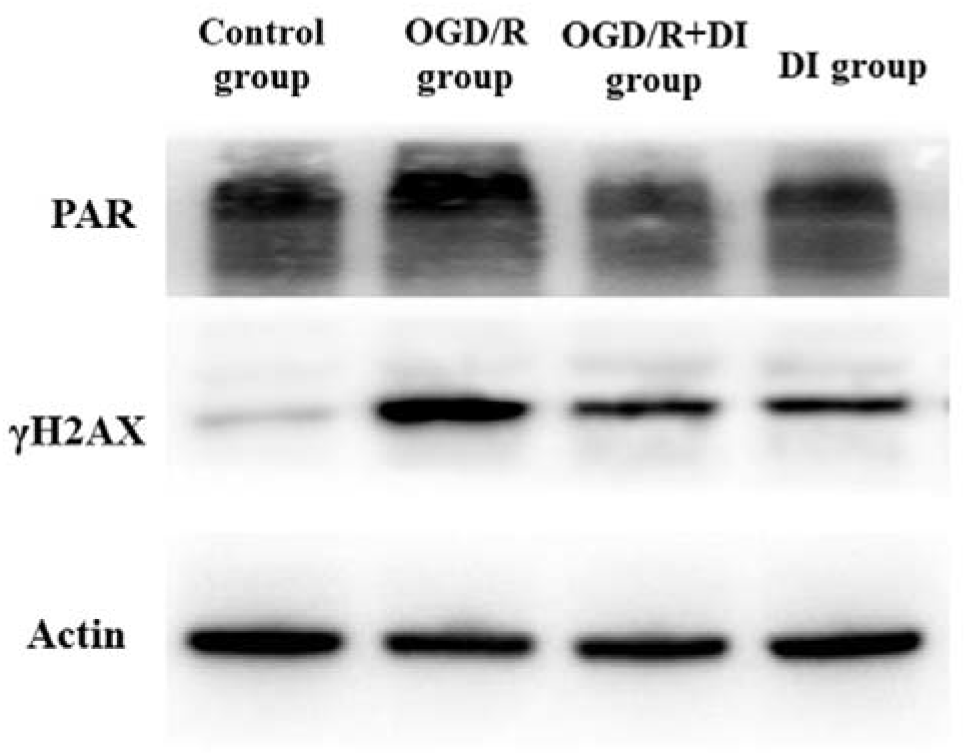
Western blotting detected the expressions of PAR and γH2AX in different groups, actin was used as the internal control. Abbreviations: PAR, Poly ADP-ribose;

**Table 4.** Comparison of the expressions PAR and γH2AX in cortical neurons in different groups 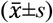.

| Group | Sample size | PAR | H2Ax |
| --- | --- | --- | --- |
| control | 3 | 0.74±0.18 | 0.40±0.13 |
| OGD/R group | 3 | 1.38±0.23<sup>①②</sup> | 1.12±0.87<sup>③④</sup> |
| DI+ OGD/R group | 3 | 0.81±0.11 | 0.64±0.13 |
| DI group | 3 | 0.97±0.29 | 0.54±0.09 |
| *F* |  | 10.26 | 23.85 |
| *P* |  | <0.01 | <0.001 |

Abbreviations: OGD/R, hypoxia/reoxygenation; DI, Dihydromyricetin; Compared with the control group, <sup>#</sup>P<0.05, <sup>##</sup>P<0.01, <sup>###</sup>P<0.001; Compared with the DI+OGD/R group, <sup>\*</sup>P<0.05, <sup>\*\*</sup>P<0.01, <sup>\*\*\*</sup>P<0.001.

## Discussion

HIE is a type of hypoxic-ischemic brain injury caused by partial or complete hypoxia of brain tissue during the perinatal period, which is one of the major causes of intellectual development, even cerebral palsy and death in newborns; the pathogenesis of HIE includes neurotoxicity, oxidative stress, and immune inflammation [19]. Many kinds of risk factors including oxidative stress and excitatory toxicity could result in neuronal death and neuronal apoptosis to lead to a series of neurological deficits [20]. Therefore, inhibition of neuronal death or reduction of neuronal apoptosis is one of the important therapeutic strategies for HIE treatment; hypoxia as one of the main ways to result in cell death and apoptosis, it could also induce the apoptosis of neurons.

Parthanatos is a mode of cell death characterized by excessive activation of PARP-1, leading to the formation of PAR oligomers, inducing AIF nuclear translocation, resulting in DNA damage and cell death [21, 22] including cerebral infarction and Parkinson’s disease [23, 24]. In fact, Ischemia and hypoxia could activate neuronal parathanatos to result in cerebral infarction, and HIE has a similar pathogenesis to cerebral infarction. In our study, we confirmed that hypoxia was induced to decrease the neuronal activity and to increase the cell death, increase the expression of _γ_H2AX and over activation of PARP-1, promote AIF nuclear translocation, these results demonstrated that hypoxia may induce parthanatos in neurons to result in HIE, which provide a potential treatment strategy for HIE.

Furthermore, we validated dihydromyricetin, which is mainly presented in grape plant and has activities including antioxidant effect, anti-apoptosis effect, anti-tumor effect, and anti-inflammatory effect [25, 26]. found that dihydromyricetin could inhibit hypoxia-induced oxidative stress, decrease the expressions of PARP-1 and _γ_H2AX, reduce the AIF nuclear translocation. Those evidences demonstrated that dihydromyricetin could reduce neuronal death and oxidative stress by regulating parathanatos, may provide some useful evidences for dihydromyricetin treatment on HIE.

## Conclusion

Dihydromyricetin alleviated the damage of hypoxia-induced mouse neurons by reducing the ROS levels and inhibiting the expressions of PAR and _γ_H2AX, which may become an ideal therapeutic drug for HIE.

## Acknowledgements

None.

